# Development and Outcomes of an International Certificate Program in Science Policy and Advocacy for STEM PhD Students and Postdocs

**DOI:** 10.1101/2023.11.26.568726

**Authors:** Adriana Bankston, Amy Ralston, Joanne Ly, Harinder Singh

**Affiliations:** Strategic Advisor, Journal of Science Policy & Governance; Graduate Student Researcher, UC Irvine; Program Director, GPS-STEM, UC Irvine; Former Program Director, GPS-STEM, UC Irvine

## Abstract

Scientists must play a significant role in enacting societal change by educating and advising policymakers on relevant policy topics. To fill the identified gap in science policy and advocacy training, during the pandemic years, we piloted and offered the online Science Policy & Advocacy for STEM Scientists Certificate Program starting in 2020. Over three cohorts, the program was focused on practical skills and concepts, networking, and career development opportunities, and providing pathways for PhD students and postdoctoral researchers to learn about and in many cases fully transition into science policy and advocacy roles and careers. Program participants were exposed to many important aspects of what it means to be involved in science policy and advocacy, and many chose to enter the field through subsequent opportunities facilitated by the program training. We sought to reduce a number of barriers to program participation, and additionally provided resources for others to develop similar programs at their university. We believe this training model is innovative and can be recapitulated, and sincerely hope to see more universities create similar programs in the US and internationally in order to serve trainees interested in science policy and advocacy and facilitate their building impactful careers in the field for years to come.

## Introduction

During the COVID 19 pandemic, we observed a significant disconnect between scientists and policy makers, which undermines our ability to address global issues in science and policy. This issue has galvanized scientists who are now more interested in science policy knowledge and careers in the field, including early career scientists (1).

Traditionally, science policy has been defined as “the set of rules, regulations, methods, practices, and guidelines under which scientific research is conducted” (2). Science and policy are interconnected and influence each other. The two fundamental concepts in policy are ‘science for policy’ and ‘policy for science.’ The first uses scientific findings to inform policy, and the second refers to policies that affect the practice of science, and how science is done.

The ways in which science has been engaged with and supporting government decision-making has evolved over the last 80 years (3). The process involves translating the different languages of scientists and policymakers back and forth, and aligning information needs with outputs. Policymakers rely on scientific advice and evidence to inform their decision-making, and the way they formulate these views takes into account many different factors within the legislative, executive, and judiciary branches of government (4). This back-and-forth interaction can be critical for how scientists inform government decision making and the interplay between the two in specific disciplines (5).

### Brief History of Science Policy

Before World War II, science was rarely in effective contact with policymaking and the political processes (3). Policymaking was generally understood to act in pursuit of “the common good,” and essentially in the best interest of the public that the government serves (6). This may occur through science advice mechanisms to governments or communicating with the public.

While policymaking may be based on scientific evidence, it’s not always clear how this knowledge enters the policymaking process, and what structured mechanisms exist for influencing policy making. Policymaking is dynamic and takes into account many different types of input, and the most effective means of connecting science with policy is also debatable.

In the book “The honest broker,” Roger Pielke describes four categories or roles of scientists in their interaction with the policy community (2). Most interest has focused on his distinction between the ‘issue advocate’ and the ‘honest broker of policy alternatives’. The issue advocate provides evidence synthesis with a specific agenda or programmatic outcome. In contrast, the honest broker provides an unbiased view of the evidence that exists to assist policymakers in making choices between different options. The other two options include the science arbiter which has a narrow focus on some questions that science can answer and does not influence the direction of policy; and pure scientists who did not engage with the policy community. Scientists may choose to engage with the policymaking process in a number of ways, and different skill sets are needed depending on their preferred method and impact sought.

### The Need for Science Policy Engagement

As we emerge from the COVID-19 pandemic, discussions needed to prevent future pandemics should be at the forefront. In order to ensure a sustainable recovery from the pandemic, it is imperative to develop effective and ethical policies to prevent future ones. This requires collaborations between scientists and policymakers in order to make real-world impact on a number of large societal issues. Therefore, scientists need to undergo specific policy training in order to understand the policy making process and the best mechanisms for engaging policy makers. This includes understanding how to allocate research budgets to identifying and sequencing viruses, predicting mutations, research on gain and loss of function studies and best practices and policies to contain the future spread of pandemics, where the development of therapies and vaccines should be at the forefront.

One way to achieve this goal includes increasing the direct and effective involvement of scientists in the policymaking process, and many individuals who work in science policy are liaisons between researchers and policymakers (7). Scientists are not only aware of the current landscape of science policy issues, but also have significant background knowledge, training, and practical advice in finding plausible solutions from gathering data and/or from their own lived experience. As experts in a number of scientific fields, scientists have an obligation and an opportunity to inform and influence the policymaking process with their expertise in an evidence-based manner. However, most scientists lack awareness and training in public policy and advocacy, which we seek to address in part with this Certificate Program.

### Current Science Policy Training Landscape for PhD Students and Postdocs

The US science, technology, engineering, and mathematics (STEM) labor force represents 23% of the total US labor force, contributing to maintaining our nation’s competitiveness in science and technology through respective employment sectors (8). Among STEM PhD students, there is a decrease in the attractiveness of faculty careers and an increase in the attractiveness of nonacademic careers (9).

PhD students became interested in careers in government towards the end of their degree as opposed to earlier (9). However, there is not enough awareness about careers in science policy and advocacy for scientists. In addition, PhD students need to be exposed to non-academic career options earlier, including those in science policy and advocacy, which have grown in interest among future scientists.

In order to pursue their career of choice, PhD students need specialized training embedded into the graduate curriculum and postdoctoral training which showcases the value of their science in society. Currently, the idea of providing tailored career and professional development training for current PhD students and postdocs is still slow or has not yet been widely adopted in academia. However, the culture is slowly beginning to change.

In 2016, National Institutes of Health (NIH) Commons Fund initiated a large career and professional development initiative for PhD students and postdocs called Broadening Experiences in Scientific Training (BEST). A total of 17 institutions in the US were awarded NIH-BEST grants, including the University of California (UC) Irvine. At UC Irvine, this led to the development of the Graduate Professional Success in Biomedical Sciences (GPS-BIOMED) program which later became GPS-STEM. The NIH-BEST programs are based on the skills assessment platform, my Individual Development Plan (myIDP). The job sectors for STEM PhDs include private industry, government, and non-profit roles. Out of the ∼20 possible career choices for PhDs in myIDP, science policy and advocacy is an area of strong career interest for scientists based on their skills, interests and values signifying the importance of investing in this training sector for academic scientists and continuing to provide opportunities (10).

Training in science policy and advocacy is currently still relatively low in universities across the country, placing many promising academic scientists at a disadvantage for this career path. While some institutions offer science policy training at the university level, the vast majority of these programs are concentrated only at a set number of institutions and do not cover all career levels. To assess the programs offered, we surveyed science policy programs from universities and organizations and obtained a list of a subset of science policy training programs in the US (**Table 1**) (references 1, 11, 12 and additional research).

**Table 1.** Currently Available Policy Training Programs. This table represents a subset of science policy training programs across the country including certificate programs to provide a flavor for the types of similar opportunities that exist, including the Certificate Program described in this publication.

| **Program Title** | **Trainees Served** | **Cost** | **Website** |
| --- | --- | --- | --- |
| University of California, Irvine Science Policy & Advocacy for STEM Scientists Certificate Program | Any affiliation | Free | https://gps-stem.grad.uci.edu/scipol/ |
| University of California, Riverside Certificate in Science Policy | Students at the institution | Free | https://sciencetopolicy.ucr.edu/ |
| University of Colorado Graduate Certificate Program in Science & Technology Policy | Students at the institution | Free | https://sciencepolicy.colorado.edu/students/ |
| University of Michigan Graduate Certificate in Science & Technology Public Policy | Students at the institution | Free | https://stpp.fordschool.umich.edu/graduate-certificate |
| Catalyzing Advocacy in Science and Engineering (CASE) Workshop | Any affiliation | Must apply for sponsorship from a participating institution | https://www.aaas.org/programs/catalyzing-advocacy-in-science-and-engineering |
| MIT "Policy for Science, Technology and Innovation" Course | Any affiliation | Free | https://www.edx.org/course/policy-for-science-technology-and-innovation |
| Science and Technology Policy Academy | Any affiliation | ~\$400-\$1000 | https://www.scitechpolicyacademy.com/ |

In order to fill this knowledge and training gap, many STEM trainees have to resort to finding opportunities on their own in science policy and advocacy. A number of graduate student and postdoctoral researchers at US universities have formed science policy affinity groups, which are composed of like-minded trainees interested in pursuing a career in science policy and advocacy with programming being organized to facilitate engagement in policy making. We compiled a subset of policy programs at different US universities to showcase a sampling of the different types of opportunities available in policy and advocacy (**Table 2**).

**Table 2.** Programs from Science Policy Groups. This table represents a subset of science policy programs from science policy groups within US universities to provide examples of trainings developed by trainees.

| Program Host | Program Type | Website |
| --- | --- | --- |
| Science Policy Group at UCSF | SciPol minicourse | https://www.spgatucsf.org/ |
| Science Policy Group at UCLA | Panel discussions | https://www.scipolucla.com/current-upcoming |
| Yale Science Diplomats | Bootcamps | https://sciencediplomats.sites.yale.edu/ |
| Science Policy Group U of Illinois Urbana-Champaign | Discussion groups | https://extension.illinois.edu/ispp |
| MIT Science Policy Initiative | Bootcamp | https://mitspi.squarespace.com/bootcamp |
| Harvard GSAS Science Policy Group | Career panels | https://projects.iq.harvard.edu/sciencepolicy/career-panels |
| Science Policy and Advocacy Group (SPAG) UNC Chapel Hill | Discussions | https://heellife.unc.edu/organization/spag |
| Penn Science Policy Group | Writing groups | https://www.pspdg.com/ |
| Science Policy Education and Advocacy Club (SPEAC) at Virginia Tech | Workshops and competitions | https://speacvt.wixsite.com/sciencepolicyvt |
| Houston Science Policy and Advocacy Group | Discussions | https://www.houstonspag.com/ |

While graduate student and postdoctoral-led programs in universities are useful for trainees to develop their own career development and can be impactful experiences, institution-organized trainings in policy and advocacy are essential for training future scientists in this country. In addition, due to the importance of evidence-based policy for achieving societal change, scientists should be educated on the policymaking process and universities should play a major role by offering short programs, workshops, bootcamps, certificate courses and for-credit academic elective courses during graduate and postdoctoral education.

## Certificate Program in Science Policy and Advocacy for STEM Scientists

### Program Development

To fill the gaps training and enhance the preparation of scientists for future careers in policy and advocacy, a collaborative initiative was developed between The University of California, Irvine (UC Irvine) Public Policy Prep (P3) program funded by [Burroughs Wellcome Fund](https://www.bwfund.org/) (part of UC Irvine’s GPS-STEM program) in conjunction with the *[Journal of Science Policy & Governance (JSPG)](http://www.sciencepolicyjournal.org/)*, the [Union of Concerned Scientists (UCS)](https://www.ucsusa.org/) and [UC Irvine’s Ridge to Reef](https://ecoevo.bio.uci.edu/graduates/ridge-to-reef/) program. The training resulted in a Certificate Program in Science Policy and Advocacy for STEM Scientists, which launched in 2020 and was open to all PhD students and postdoctoral fellows from anywhere in the world, without restrictions on career stage, policy knowledge or citizenship status (13).

Over the course of three years, this program was organized in conjunction with three founding partners: *Journal of Science Policy & Governance (JSPG)*, UC Irvine’s Ridge to Reef, and the organization Union of Concerned Scientists (UCS) which were instrumental in program concept development and recruitment of PhD students and postdocs, as well as in bringing expertise to the program itself through innovative speakers, workshop organizers and facilitators with experience knowledge in science policy and advocacy. Additionally, the program was financially supported in part by [STEMPeers](https://stempeers.org/) and [UCS Science for Public Good Fund](https://www.ucsusa.org/resources/science-public-good-fund). In 2023, STEMPeers, [Sigma Xi](https://www.sigmaxi.org/), and [UC Irvine Science Policy and Advocacy Network (SPAN)](https://www.linkedin.com/in/uci-span-a5a0a921a/) contributed speakers and brought new perspectives to the program (14).

### Program Goals and Specific Outcomes

The main goals of the program were to provide awareness about careers in science policy and advocacy to PhD students and postdocs, and offer specialized training and skill set building to develop policy leaders of the future. Specifically, the program provided training in written and oral communication in science policy and practical experiences, as well as opportunities for networking with professionals.

#### Program Outcomes

- Build expertise in giving an elevator pitch to policymakers
- Grow network among peers and other science policy experts
- Perform informational interviews with science policy professionals
- Learn fundamentals of science policy and advocacy from various resources

***Core Competencies*** for the programs include: critical thinking, problem solving, cross-discipline applications, community building, and building diversity and inclusivity.

***Skills*** that participants learned include: oral and written communication, teamwork, leadership, empathy, persuasion, and ethical reasoning.

### Program Structure and Methods

Given the program was started during the pandemic, we took advantage of the virtual space and enrolled a large number of trainees, both within the US and internationally, for three years in a row (2020, 2021, 2023). The graduation rate was high all three years and participants received certificates of course completion.

The original concept for the program was designed in 2020 and composed of a total of 9 modules plus 1 writing training module and 1 elevator pitch competition (**Table 3**). Each learning module of the program had a pre-reading assignment, and several different types of homework after each session for which reviewer feedback was solicited to improve oral and written skills in multiple rounds. The final session was an elevator pitch competition with cash prizes. Subsequent iterations of the program were modified in a flipped-classroom format in 2021, and a hybrid between the two formats in 2023 with common elements throughout the three cohorts (**Supplementary Figure 2**). Data presented in the publication results are either from two or all three years cohorts.

**Table 3.** Certificate Program Modules, Description and Assignments. Links in this table contain YouTube video content from the 2020 program, which was the original program concept. Additional video playlists from the 2021 and 2023 program versions can be found in Supplementary Figure 2.

| Module | Module Title | Module Description | Assignments Following Listed Class Date |
| --- | --- | --- | --- |
| 1 | **Introduction to Science Policy & Advocacy**<br>Module 1 Playlist | - Explains the fundamentals of science policy and advocacy, differentiating between ‘science for policy’ and ‘policy for science.<br>- Explores general approaches for researching and pitching on science policy and advocacy topics and exemplifies responsible communication skills for advocating on Capitol Hill. | - Listen to and summarize science policy podcast episode<br>- Connect with podcast guest on LinkedIn<br>- Brainstorm topics for science policy pitch |
| 2 | **Course Orientation, Information, Q&A and Scientific Research Policy**<br>Module 2 Playlist | - Exposes participants to a few topics in scientific research policy with broad societal relevance, providing a flavor for research advocacy. | - Begin drafting oral elevator pitch outline based on your topic of choice<br>- Listen to podcast episode for education and workforce development<br>- Come up with a question to ask during next class |
| Writing | **Writing module**<br>Writing Module Playlist | - Describe how policy writing differs from academic writing, provide a few examples of different types of policy writing, breakdown elements of such a write-up and begin to craft a written policy piece of your choice. | - Continue drafting your policy pitch<br>- Begin drafting you short policy write-up and receive feedback |
| 3 | **STEM Education and Workforce Development**<br>Module 3 Playlist | - Speakers describe ways to improve the research enterprise by increasing equity and inclusion and engaging in professional development opportunities for transitioning into non-academic careers. | - Brainstorm topics to include in policy write-up<br>- Continue to work on policy write-up |
| 4 | **Effective Advocacy Strategies for Policymakers**<br>Module 4 Playlist | - Discusses what advocacy means, covers ways of interacting with policymakers at the federal and state level, and provides practical ways to advocate through a role-playing exercise. | - Pick a policymaker to give your pitch to<br>- Design and submit a one pager about your policy issue detailing how you would advocate for the issue on the Hill, and how you would frame the issue for their office with<br>details on this person and their policy interests<br>- Submit elevator pitch to fit that particular policymaker |
| 5 | **AAAS STPF for Transitioning into Science Policy & Advocacy Careers STPF panel**<br>Module 5 Playlist | - The AAAS Science and Technology Policy Fellowships (STPF) program provides an innovative and robust way to train the next generation of science policy leaders.<br>- Speakers discussed experiences with the STPF program and answer questions about the application process. | - Do an informational interview with a fellow (STPF)<br>- Upload google doc with answers to provided questions from informational interview |
| 6 | **State and Federal Policy Fellowships for Transitioning into Science Policy & Advocacy Careers**<br>Module 6 Playlist | - In addition to AAAS STPF, other state and local science policy fellowships can facilitate transitions into policy careers.<br>- Speakers in this module discussed their experiences with a variety of fellowships, both at the state and national levels, and how these fellowships advanced their careers. | - Do an informational interview with someone a fellow other than STPF (state and federal)<br>- Upload google doc with answers to provided questions from informational interview |
| 7 | **Effective Strategies for Targeted Policy & Advocacy Engagement**<br>Module 7 Playlist | - Provided an overview of how science communication strategies can be applied to science advocacy and policy.<br>- Participants heard from a UCS policy expert about ways to build relationships with policymakers as a way of informing and influencing the policy making process.<br>- The trainers introduced power mapping as a tool for better understanding policymakers so you can craft your most effective messages to inform or persuade them. | - Submit rough draft of power mapping plan<br>- Submit revised elevator pitch<br>- Sign up for elevator pitch practice session |
| Training | **Elevator pitch practice session and feedback** | - This session served as a practice for the elevator pitch in front of organizers and other experts, with judging and opportunities to advance to the final competition. | - Submit elevator pitch<br>- Continue working on policy write-up |
| 8 | **Leveraging Science Policy & Advocacy for Non-academic and Industry Careers**<br>Module 8 Playlist | - Discussed how science policy awareness and analysis training can be applied to careers in non-academic and industry sectors.<br>- Highlighted the repertoire of transferable skills to consider when pursuing non-academic and industry careers.<br>- Individuals from a range of industry and government careers provided perspectives on utilizing science policy skills for career success. | - Continue working on policy write-up<br>- Do an informational interview with an industry professional<br>- Upload google doc with answers to provided questions from informational interview |
| 9 | **Science Policy & Advocacy Engagement at the Local Level**<br>Module 9 Playlist | - Experts in this module work in various sectors of society, and they will discuss how science policy and advocacy can improve scientific literacy and public engagement. | - Identify a public engagement project of interest<br>- Craft a short pitch idea of how you would engage locally on your topic<br>- Submit policy write-up |
| Competition | **Elevator Pitch Competition**<br>Competition Playlist | - Learning how to pitch your research to policymakers is critical for science policy. Selected students participated in the final competition. | N/A |

### Program Activities

- Listening to [SciPol SoundBites (JSPG)](https://www.sciencepolicyjournal.org/podcast), GPS-STEM Radio-Science Policy Podcast and other podcasts to learn about science policy careers
- Constructing an oral policy elevator pitch for policymakers
- Writing op-ed or policy memo on policy topic of choice
- Designing advocacy one pager for policymakers
- Crafting power mapping plan on policy topic of interest for policymakers
- Participating in policy elevator pitch competition

### Program Deliverables

- Podcast summary from SciPol SoundBites (JSPG), GPS-STEM Radio - Science Policy Podcast or other podcasts
- Elevator pitch video recording submission
- Policy writing piece (policy memo, op-ed)
- One pager talking points and design for policymakers
- Informational interview Q&A responses
- Power mapping plan for policymakers
- Public engagement plan for local/state engagement
- Participate in policy pitch competition (if selected)
- Interview policy professionals on GPS-STEM Radio Science Policy podcast (winners)

## Results and Discussion

### Program Participant Data

Program participants were from a number of US and international institutions, showing a broad interest in policy and advocacy from trainees all over the world and from a number of sectors, with the vast majority from academia and a large number of universities but also from industry, government, and nonprofits (**Table 4**).

**Table 4.** Institutions of Program Participants. These data constitute the institutions from which trainees who enrolled in the program initially come from and their affiliations at the time of taking the class or at the time of graduation, encompassing PhD students, post-PhD, and pre-PhD trainees.

| USA Locations |  |  |
| --- | --- | --- |
| State | Type | Name |
| AK | University | University of Alaska Fairbanks |
| AL | University | Auburn University |
| AL | University | University of Alabama at Birmingham |
| AZ | University | Arizona State University |
| AZ | University | University of Arizona |
| CA | University | Harvey Mudd College |
| CA | Institute | Scripps Research Institute |
| CA | University | UC Berkeley |
| CA | University | UC Davis |
| CA | University | UC Irvine |
| CA | University | UC Los Angeles |
| CA | University | UC San Diego |
| CA | University | Stanford University |
| CO | Institute | NOAA Earth System Research Laboratories (ESRL) |
| CO | University | University of Colorado-Anschutz |
| CO | University | Colorado State University |
| CT | University | Yale University |
| DC | Agency | United States Agency for International Development (USAID) |
| DC | University | George Washington University |
| DC | University | Georgetown University |
| DC | Think tank | International Council on Clean Transportation |
| DC | Institute | Children's National Hospital |
| DE | University | Kesmonds International University |
| FL | University | Florida International University |
| FL | University | University of Florida |
| GA | University | Emory University |
| GA | University | Georgia Institute of Technology |
| GA | University | Morehouse School of Medicine |
| GA | University | University of Georgia |
| IA | University | Iowa State University |
| IL | University | Illinois State University |
| IL | University | Northwestern University |
| IL | University | University of Illinois Urbana-Champaign |
| IL | University | University of Chicago |
| IL | University | University of Illinois at Chicago |
| IN | University | Indiana University School of Medicine |
| IN | University | Purdue University |
| LA | University | Louisiana State University |
| LA | University | University at Louisiana at Lafayette |
| KS | University | Kansas State University |
| MA | Organization | ACLU of Massachusetts |
| MA | Institute | Broad Institute (MIT and Harvard) |
| MA | University | Harvard University |
| MA | University | Massachusetts General Hospital/McLean |
| MA | Institute | Massachusetts Institute of Technology |
| MA | Institute | Ragon Institute of Mass General (MIT and Harvard) |
| MA | University | Tufts University |
| MA | University | University of Massachusetts |
| MD | Institute | National Institute of Health (several institutes) |
| MD | University | Johns Hopkins University |
| MD | Agency | Food and Drug Administration (FDA) |
| MD | University | University of Maryland Baltimore |
| MD | University | University of Maryland |
| MD | University | Uniformed Services University |
| MD | University | Johns Hopkins University |
| MI | University | Michigan State University |
| MI | University | University of Michigan |
| MN | University | Mayo Clinic |
| MN | University | University of Minnesota |
| MO | University | University of Missouri-Kansas City |
| MO | University | Washington University in St. Louis |
| MS | University | Mississippi State University |
| NC | University | Duke University |
| NC | University | East Carolina University |
| NC | University | UNC Chapel Hill |
| NC | University | UNC Greensboro |
| NC | University | North Carolina A&T State University |
| NC | University | North Carolina State University |
| NC | University | Wake Forest University School of Medicine |
| NE | University | University of Nebraska- Lincoln |
| NH | University | Dartmouth College |
| NM | University | University of New Mexico |
| NY | Institute | Regeneron Pharmaceuticals |
| NY | University | SUNY Downstate |
| NY | University | SUNY Stony Brook University |
| NY | University | Albany Medical Center |
| NY | University | Albert Einstein College of Medicine |
| NY | Institute | Cold Spring Harbor Laboratory |
| NY | University | University of Rochester Medical Center |
| NY | University | Columbia University |
| NY | University | Cornell University |
| NY | University | Mount Sinai School of Medicine |
| NY | University | Memorial Sloan Kettering Cancer Center |
| NY | Institute | NASA Goddard Institute for Space Studies, Columbia University |
| NY | Institute | New York City Department of Health and Mental Hygiene |
| NY | Institute | New York Genome Center |
| NY | University | Rockefeller University |
| NY | University | University at Buffalo |
| NJ | University | Rutgers University |
| NJ | University | Princeton University |
| OH | Institute | Cleveland Clinic |
| OH | University | The Ohio State University |
| OH | University | The College of Wooster |
| OH | University | University of Cincinnati |
| OR | University | Oregon State University |
| OR | University | University of Oregon |
| PA | Agency | PRECISIONscientia |
| PA | University | Thomas Jefferson university |
| PA | University | University of Pennsylvania' |
| PA | University | University of Pittsburgh |
| RI | University | Brown University |
| SC | University | Clemson University |
| SD | University | University of South Dakota |
| TN | University | University of Tennessee, Knoxville |
| TN | Institute | USDA-Agricultural Research Service |
| TN | University | Vanderbilt University |
| TX | University | Baylor College of Medicine |
| TX | University | Rice University |
| TX | University | Texas A&M University |
| TX | University | University of Texas at Austin |
| TX | University | University of Texas MD Anderson Cancer Center |
| UT | University | University of Utah |
| VA | University | George Mason University |
| VA | Institute | Research!America |
| VA | University | Virginia Tech |
| VA | Agency | US Department of Defense |
| VA | University | University of Virginia |
| VA | Institute | St. Jude Children's Research Hospital |
| VT | University | University of Vermont |
| WA | University | University of Washington |
| WI | University | University of Wisconsin - Madison |
| **International (non-US) Locations** |  |  |
| **Country** | **Type** | **Name** |
| Africa | Company | SynBio Africa |
| Bangladesh | University | Jahangirnagar University |
| Brazil | University | UNICAMP Universidade Estadual de Campinas |
| Brazil | University | University of Sao Paulo |
| Burundi | Institute | Prince Régent Charles Hospital |
| Canada | University | University of Guelph |
| Canada | University | University of New Brunswick |
| Canada | University | University of Toronto |
| Canada | University | Dalhousie University |
| Canada | University | University of Alberta |
| Canada | University | Memorial University of Newfoundland |
| Canada | University | McGill University |
| Canada | University | McMaster University |
| Canada | University | Queen's University |
| Canada | University | Université de Montréal |
| Canada | Institute | Institut national de la recherche scientifique (INRS) |
| Costa Rica | University | University of Costa Rica |
| France | University | Sorbonne University |
| France | Institute | Institut NeuroMyoGène |
| Germany | University | Justus Liebig University Giessen |
| Germany | Business | Trilogy Writing & Consulting |
| Germany | University | Justus Liebig University Giessen |
| Germany | University | Max Planck Institute for Biophysical Chemistry |
| Jamaica | University | University of the West Indies |
| Japan | University | University of Hyogo |
| India | University | St.Xavier's College (Autonomous), Kolkata |
| India | University | Virginia Tech India |
| India | University | St. Petersburg College |
| India | Institute | Indian Institute of Science |
| India | University | Sambalpur University |
| India | Institute | Rajiv Gandhi Centre for Biotechnology |
| India | Institute | All India Institute of Medical Sciences |
| India | Institute | CSIR-Institute of Genomics & Integrative Biology (IGIB) |
| India | Institute | CSIR-National Institute of Science Communication and Policy Research (CSIR-NIScPR) |
| India | Institute | Indian Institute of Science |
| India | University | Christian Medical College |
| India | Organization | National Council of Educational Research and Training |
| India | University | Daffodil International University |
| India | Organization | Department of Science and Technology, Govt. of India |
| India | University | Jawaharlal Nehru University |
| Italy | University | Università del Piemonte Orientale |
| Italy | University | University of Bologna |
| Jakarta | Institute | Indonesian Institute of Sciences |
| Japan | University | Tokyo University of Science |
| Jordan | University | Jordan University of Science and Technology |
| Kenya | University | Pan African University Institute for Basic Sciences, Technology, and Innovation (PAUSTI) |
| Kenya | University | Jomo Kenyatta University Of Agriculture And Technology |
| Mauritius | University | University of Technology, Mauritius |
| Malaysia | University | Monash University |
| Malawi | University | Kamuzu University of Health Sciences (KUHeS) |
| Mexico | University | National Autonomous University of Mexico |
| Mexico | Institute | Colegio de Posgraduados |
| Mexico | University | ENCB - Escuela Nacional de Ciencias Biológicas Unidad Santo Tomás IPN |
| Mexico | Business | MY World Mexico |
| Mexico | Institute | Cinvestav |
| Mexico | Institute | Instituto Politécnico Nacional |
| Mumbai | University | University of Mumbai |
| Nepal | University | Kathmandu University School of Arts |
| Netherlands | University | Radboud University Medical Center |
| Nigeria | University | Bayero University, Kano |
| Nigeria | Institute | Pharmacy Council of Nigeria |
| Nigeria | University | University of Nigeria, Nsukka |
| Nigeria | University | Kaduna State University |
| New Zealand | University | University of Otago |
| Norway | University | University of Bergen |
| Panama | University | Universidad de Panamá |
| Pakistan | University | Shaheed Benazir Bhutto University |
| Pakistan | Institute | The Karachi Institute of Biotechnology and Genetic Engineering (KIBGE) |
| Pakistan | University | University of Health Sciences Lahore |
| Peru | University | Universidad Nacional Autónoma de Alto Amazonas |
| Philippines | Institute | UP Manila National Telehealth Center |
| Philippines | University | University of the Philippines Manila |
| Puerto Rico | University | University of Puerto Rico |
| Rwanda | University | University of Rwanda |
| Saudi Arabia | University | Qassim University |
| Scotland | University | The University of Edinburgh |
| Spain | University | Universitat Autònoma de Barcelona |
| South Africa | University | Durban University of Technology |
| South Africa | University | University of Kwazulu-Natal |
| South Asia | University | Tribhuvan University |
| Sri Lanka | Institute | National Science and Technology Commission of Sri Lanka (NASTEC) |
| Sri Lanka | Foundation | National Science Foundation of Sri Lanka |
| Switzerland | University | ETH Zurich and the École Polytechnique Fédérale de Lausanne (EPFL) |
| Turkey | University | Istanbul Technical University |
| Uganda | University | Makerere University |
| United Kingdom | University | Newcastle University |
| United Kingdom | University | University College London |
| United Kingdom | University | University of Glasgow |
| United Kingdom | University | Lancaster University |
| Zimbabwe | University | University of Zimbabwe |

Across the three cohorts, an increasing number of trainees registered from 2020 to 2023, with selection increasing in 2021 and 2023 with the application process, with relatively high retention and graduation rates (**Supplementary Table 1**). During peak pandemic times, some trainees were unable to complete the program due to personal reasons including either themselves or their families getting sick from COVID-19. A few were unable to complete the minimum requirements for the program and were not offered the certificate. However, overall interest in policy and advocacy remained high and the program served trainees from multiple countries without restrictions.

Participants who enrolled in the program initially were majority PhD students (66.1%), followed by post-PhD (21.1%), pre-PhD (9.4%) and other/unspecified (3.4%) (**Figure 1A**). These data show a strong interest from graduate students to enroll in the program, but also from other career stages. A higher number of pre-PhD students enrolled in 2023 which may indicate developing an interest in the field earlier in their training (**Supplementary Table 2**) and this could shift the paradigm towards how the future of the field looks and who is able to contribute.

**Figure 1.**
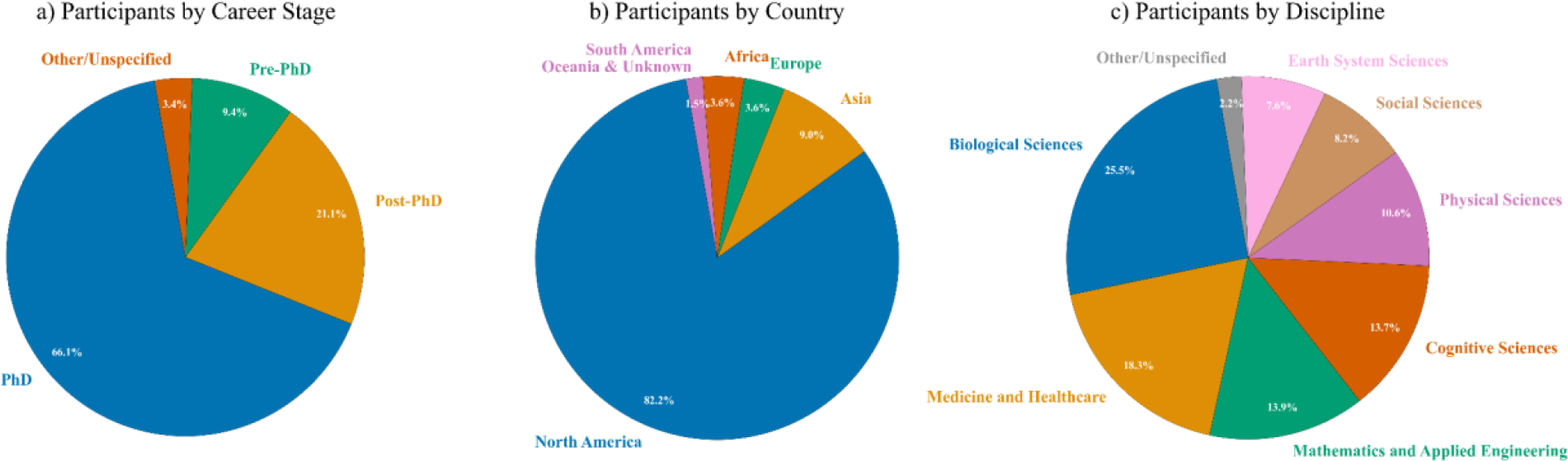
Participant Data. These data constitute the percentage of trainees who enrolled in the program initially, by career stage (A, all 3 cohorts), by country (B, 2020 and 2023) and by discipline (C, all 3 cohorts). For (A), career stages included PhD students, post-PhD, and pre-PhD. The post-PhD career stages included postdoc, research staff, project scientist, specialist. The pre-PhD career stages included undergraduate and masters students. For (B), this included geographical location when taking the course based on affiliation. For (C), disciplines were broadly defined due to the variability of participant entries and interdisciplinary nature of their studies. For the publication, we assigned disciplines based on broad categories around specific scientific interests of participants.

In terms of where program participants were located or studied while enrolled in the program, the majority were from North America (82.2%), followed by Asia (9%), Europe (3.6%) tied with Africa (3.6%), followed by South America, Oceania and a few unknown combined (1.5%) (**Figure 1B**). These data likely reflect the best time zones for the participants given the majority US, including potentially the largest number of applicants being from the US. However, several trainees joined from other countries often at odd hours and were among the graduating class, going on to pursue other policy activities, signifying a growing interest in the field around the world.

A detailed breakdown of the locations of program participants showed mainly US in 2020 and 2021, with an elevation overtime towards more non-US locations and coming close to equal in numbers in 2023 based on the initial affiliation listed when enrolled (**Supplementary Table 3**). These data suggest an increasing interest in science policy from trainees located in countries other than the US, potentially changing the landscape of the field in the future in several ways.

Program participants also came from a wide range of disciplines, defined based on broad categories, with biological sciences being the highest (25.5%), followed by medicine and healthcare (18.3%), followed by mathematics and applied engineering (13.9%) tied closely with cognitive sciences (13.7%), followed by physical sciences (10.6%), social sciences (8.2%), earth system sciences (7.6%) and other/unspecified (2.2%) (**Figure 1C**). These data indicate the broad interest in policy training from trainees in a wide variety of disciplines across the three cohorts.

### Program Satisfaction Data

In addition to tallying participant data, we asked participants about satisfaction with individual modules at the completion of the program. The data across modules was very consistent among the three cohorts. Given the number of modules, for the purposes of this publication, we elected to show results from the modules related to several deliverables that are critical for success in science policy careers, with similar results for all modules.

Satisfaction was largely rated ‘very good’ or ‘excellent’ among the three cohorts when it comes to the one pager module [Module 4 in Table 3] (**Figure 2A**), power mapping module [Module 7 in Table 3] (**Figure 2B**), writing module [Writing module in Table 3] (**Figure 2C**), elevator pitch module [Training module in Table 3] (**Figure 2D**) and fellowship panel modules [Modules 5/6 in Table 3] (**Figure 2E**). These data indicate a positive outcome in exposing participants to these important skill building sessions and the fact that they mastered these skills and concepts to a high enough level in order to obtain the certificate following reviewer feedback and completion of all assignments to a satisfactory degree.

**Figure 2.**
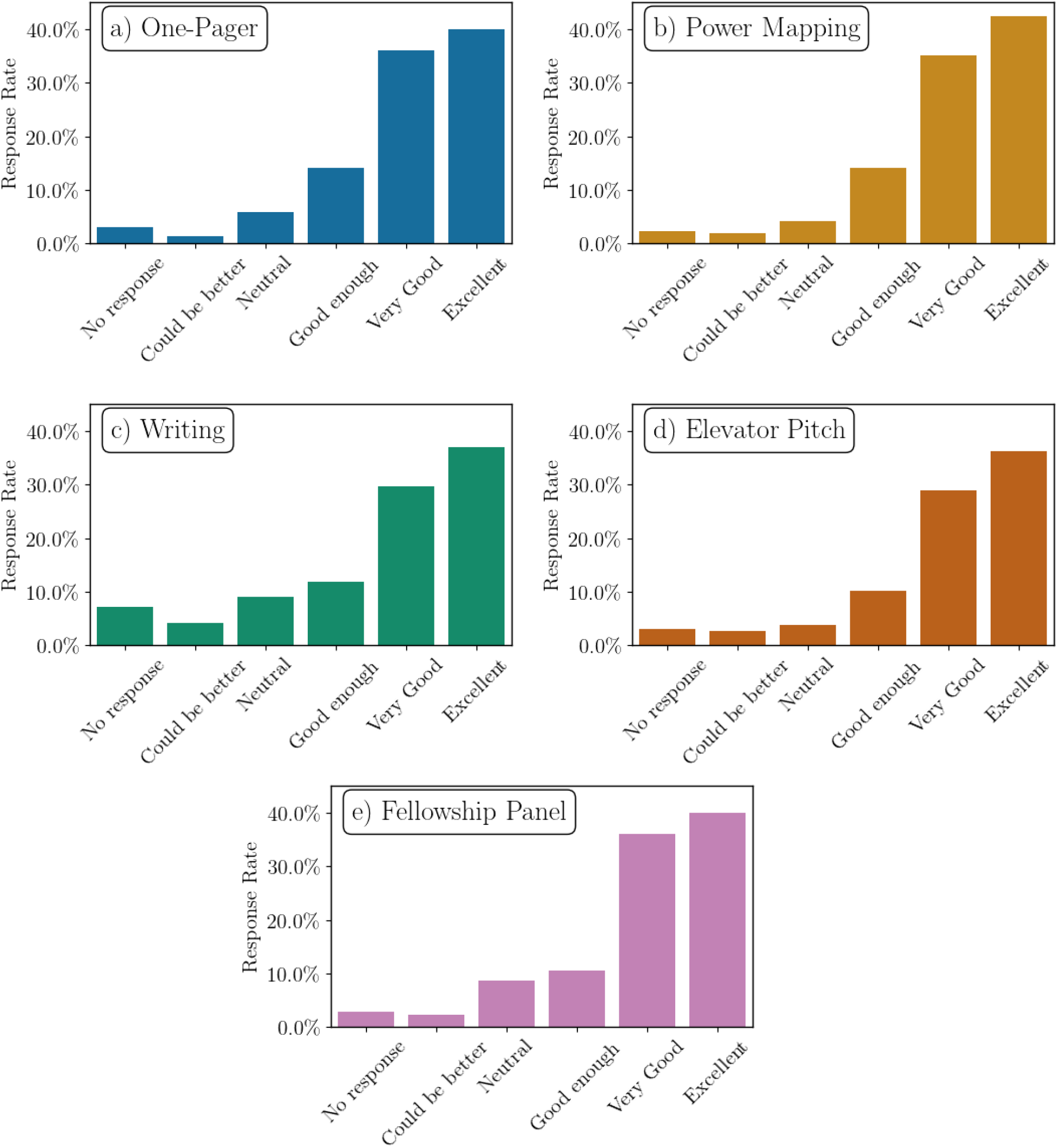
Module Satisfaction Data. These data constitute the average percentage of responses from trainees from all three cohorts who completed the program and their satisfaction with the one pager module (A), power mapping module (B), writing module (C), elevator pitch module (D) and fellowship panel modules (E) which were included in the post-program survey.

Next, we conducted surveys at the conclusion of each year’s program cohort to assess the program overall satisfaction from program graduates. Participant satisfaction overall was largely positive, with the majority either somewhat agreed or strongly agreed with survey statements in each box related to policy knowledge (**Figure 3A**) (statement: “*The course was very comprehensive, full of essential policy knowledge and essential aspects”*) and field knowledge (**Figure 3B**) (statement: “*Now I have a much better understanding of the science policy and advocacy”*). The same result was obtained when asked about whether they recommended the program (**Figure 3C**) (statement: “*This course is recommended for all STEM scientists interested in science policy as a future career”*). These data indicate the value that the participants found in the program and how it can help support them further in pursuing careers in science policy and advocacy.

**Figure 3.**
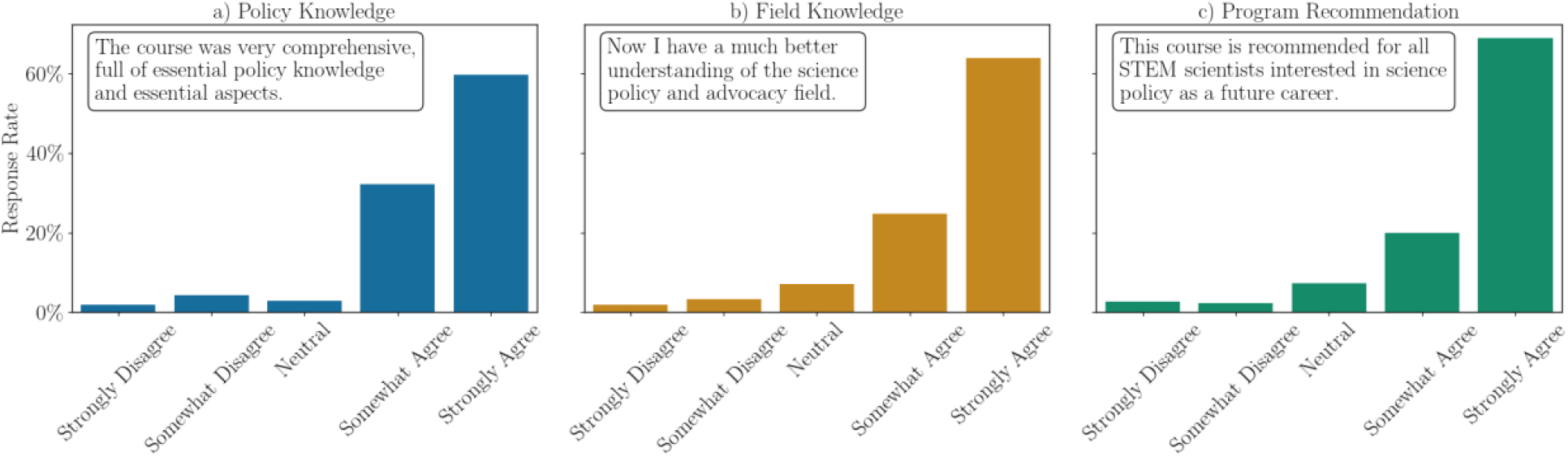
Overall Satisfaction Data. These data constitute the average percentage of responses from trainees from all three cohorts who completed the program and their overall program satisfaction when assessing policy knowledge (A), field knowledge (B) and the desire to provide a program recommendation (C) by responding to the prompts in each of the boxes which were included in the post-program survey. For (A), the statement prompt was that “*The course was very comprehensive, full of essential policy knowledge and essential aspects.”* For (B), the statement prompt was that “*Now I have a much better understanding of the science policy and advocacy.* For (C), the statement prompt was that “*This course is recommended for all STEM scientists interested in science policy as a future career.”*

### Measures of Program Success and Outcomes

To hear more about the experiences of participants in their own words, we also collected quotes to assess how the program helped them in their next steps or career trajectory in policy and advocacy. For many trainees this program kickstarted their careers in policy, while others became more confident in their decision to follow this career path after completing the program while having only considered it before.

#### Program Exposure and Promotion

Many trainees had a number of successes, some of which we presented in a poster at the National Postdoctoral Association annual meeting in 2022 (Poster slide; [Poster video](https://www.youtube.com/playlist?list=PLxh_oLbA4CUGWn6Xf6wWH9StmBUTQ81pY)) (**Supplementary Figure 1**). We also presented data about the program at several other conferences including a [FASEB conference](https://www.youtube.com/playlist?list=PLxh_oLbA4CUGWn6Xf6wWH9StmBUTQ81pY), the Graduate Career Consortium Meeting, the AAMC GREAT Meeting and the AAAS Conference, which served to raise awareness among the academic community and university staff who are working on training programs and science policy and advocacy spaces to include these programs in future offerings for PhD students and postdocs. The program was also discussed in blog posts including in [ECRlife](https://ecrlife.org/higher-education-re-imagined-goals-societal-impacts-and-pandemic-effects/) and referenced in policy [publications](https://ecrlife.org/higher-education-re-imagined-goals-societal-impacts-and-pandemic-effects/) for contributions to the science policy landscape.

#### Program Testimonials

Program graduates pursued different types of outcomes signifying their continued interest in the field upon graduation. Some of these include fellowships, publications, and policy roles as a few representative categories, along with quotes provided by participants. These data show the variety of outcomes which the participants had from and closely following involvement in their program cohort (such as being accepted to a policy fellowship, publishing a policy written piece etc.), whether they transitioned fully into policy careers or stayed in research and utilized this experience to learn about the field. These quotes show their commitment to completing the program, their enjoyment of the experience and the program contribution to their future career steps, including those from other countries who found it extremely valuable and enrolled at unusual hours (notable quote “*This course was so gripping that I woke up at 3 AM (IST) for Zoom classes,*” **Table 5**).

**Table 5.** Certificate Program Alumni Outcomes and Testimonials from the Various Cohorts. This table contains affiliations of program participants, post program accomplishments, subsequent positions, and participant quotes, in addition to quotes signifying their commitment to completing the program, their enjoyment of the experience and the program contribution to their future career steps.

| Example | Name | Affiliation at the Time of the Program | Post Program Accomplishment and Position | Quote Highlights |
| --- | --- | --- | --- | --- |
| 1 | Chrystelle Vilfranc | PhD student, University of Cincinnati | **FELLOWSHIP**<br>2021-2022 AAAS Mass Media Fellow, The Indianapolis Star. She is currently a postdoc at Columbia University. | - The science policy and advocacy course was incredibly helpful for giving me an in-depth introduction to science policy and advocacy, as well as **tangible skills that I could apply to my career and professional development.**<br>- I learned a lot about writing and advocating for policies. We were also provided a great introduction to a few individuals' experiences getting involved in the policy field. We were also given several tips and tricks for getting into policy careers. More personally, the **writing assignments helped me further stretch my writing muscles and the feedback I received was incredibly helpful.**<br>- I tweaked our public engagement **assignment, incorporating the feedback I got from the class, to apply and secure a 2021 AAAS Mass Media Fellowship.** I also plan to use the feedback from my op-ed to edit and submit it to a newspaper soon.<br>- Overall, this experience was great for my professional development. It has broadened my idea of how I can engage in policy, in some form, no matter what career I pursue. |
| 2 | Jacklyn Brennan | Postdoc, George Washington University | **FELLOWSHIP**<br>2011-2022 AAAS/ASME Congressional fellow. She is currently an AAAS fellow at USAID. | - I highly recommend this program to any grad students or postdocs interested in science policy and advocacy! I took this certificate course in 2020 and feel like it truly **gave me the knowledge and skills I needed to apply for (and successfully land!) two science policy fellowships in 2021 and 2022.** |
| 3 | Rebecca Voglewede | IRTA postdoc, National Institutes of Health | **FELLOWSHIP**<br>Selected for Rapid Response STPF Cohort in AI in 2023 | - I wholeheartedly recommend UC Irvine's Science Policy and Advocacy Certificate Program to scientists passionate about shaping the future of science and policy!<br>- **The program equipped me with the essential knowledge and skills needed to bridge the gap between science and decision-making and connected me with a wide network of peers and mentors in the science policy and advocacy world.**<br>- The program directors' care for each of the course participants is obvious and I felt incredibly supported as I pursued the next steps in my career. |
| 4 | Komal Syed | Candidate in Materials Science & Engineering, UC Irvine | **FELLOWSHIP**<br>2021-2022 Mirzayan Fellow<br>[program mentioned in fellowship bio].<br>She is currently a program officer at the NASEM GUIRR roundtable. | - This has been the most eye-opening course when it comes to **exposure to non-traditional STEM PhD careers.**<br>- I had **opportunities to network** with science policy fellows and peers of similar interests, was able to **enhance my written and oral communication skills in policy** and learned effective strategies for scientists to communicate with policymakers.<br>- **It is only because of this course that I was able to prepare to be a 2021 Mirzayan fellow at the National Academies.** |
| 5 | Ravindra Palavalli-Nettimi | Postdoctoral research associate, Florida International University | **PUBLICATION**<br>Class op-ed was published in JSPG.<br>He is currently a project specialist at the University of Tokyo. | - I was stuck in India's strict lockdown during the first wave of the Covid-19 pandemic in 2020.<br>- I saw an ad about this course which excited me because international students/scholars have very few opportunities to participate in policy-related internships in the US **This course was so gripping that I woke up at 3 AM (IST) for Zoom classes.**<br>- The course gave me an overview of policy issues and careers and motivated me to hone my skills further.<br>- By the end of this course, **I went from not knowing what science policy entails to publishing an Op-Ed in the Journal of Science Policy and Governance—all thanks to the course instructors and organizers.** I recommend this course to anyone interested in science and policy. |
| 6 | Alejandra Villegas | PhD student, University of Georgia | **EDITOR**<br>Associate editor for JSPG starting in 2021. She is currently still a graduate research assistant at University of Georgia. | - This course was hands-down the best use of my time while experiencing a pandemic. It was my first policy course, and an amazing hands-on introduction. **I walked away feeling confident about approaching informational interviews (which are crucial for any career transition), policy writing, and meetings with policymakers.**<br>- More importantly, **I was able to start building a SciPol network of people I can now reach out to for additional guidance and/or collaborations!**<br>- Since completing the science policy and advocacy for STEM scientists certificate program, I have tried my hand at 2 policy papers, met with several of my local and state elected officials, and **joined the Journal of Science Policy and Governance as an Associate Editor.** In this role, I hope to apply what I learned from this program while becoming a better science policy writer. |
| 7 | Mallory Smith | Graduate research assistant, University of Kansas | **MANAGER**<br>She is currently a Science Policy Manager at ASBMB. | - As a former participant, the **activities and certificates were IMPACTFUL when carving my path into science policy.** |
| 8 | Maksim Dolmat | Graduate student, University of Alabama | **POSTDOC**<br>Postdoctoral scholar, UC San Diego | - The program has been an absolutely incredible experience, and I've cherished every moment of it. **The program's resources, modules, networking opportunities, feedback, and shared knowledge have been nothing short of phenomenal.** It's truly remarkable how the team managed to bring it all together.<br>- I am immensely proud to have reached the finals and to have had the chance to present my pitch.<br>- **Overall, it has been an exceptionally rewarding experience, and I am sincerely thankful for being selected for this program.** I've had the opportunity to learn from some of the best in the field. |

#### Examples of Program Outcomes

Whereas few publications exist which have significantly evaluated policy programs (16), we evaluated the success based on outcomes following the program to advance the training and career development of participants in policy and advocacy.

> ***Fellowships:*** Following the program, participants were selected for competitive programs such as the AAAS fellowship (including the 2023 AAAS STPF cohort in AI) [Table 5, examples 1-3] and the Mirzayan fellowship [Table 5, example 4]. Other examples include that [Hammed Tukur](https://www.linkedin.com/in/tukurha/overlay/about-this-profile/) was selected for the Newcastle University Policy Academy Fellows Programme, showing some of the international impact.
>
> ***Publications:*** Participants published their written class assignments in *JSPG* [Table 5, example 5]. Other examples include: 1) [Marco M. Grande](https://www.linkedin.com/in/marcogrande93/), a research fellow at the University of Bologna, Department of Agricultural and Food Sciences, had his [class op-ed published in *JSPG*](https://www.sciencepolicyjournal.org/article_1038126_jspg200104.html) in March 2022; 2) [Nathan Burns](https://www.sciencepolicyjournal.org/article_1038126_jspg200104.html), a doctoral candidate at the University of Utah in the Department of Human Genetics, had his class op-ed [published in a local newspaper](https://www.sltrib.com/opinion/commentary/2023/10/04/nathan-g-burns-wic-funding-is/); 3) Pankaj Bhambhani, a science communicator, had his class op-ed published by an [India-based news site](https://science.thewire.in/the-sciences/opinion-time-to-rethink-age-restriction-jrf/).
>
> ***Policy Roles:*** Participants took on different roles during or after participating in the program, including associate editor with *JSPG* [Table 5, example 6] and science policy manager [Table 5, example 7]. Others chose to stay in research and apply this knowledge to their postdoc position [Table 5, example 8]. Other examples include [Ojas Deshpande](https://www.linkedin.com/in/ojasdeshpande/) who became a scientific policy officer at the BarcelonaBeta Brain Research Center (BBRC).
>
> ***Blogs and Programs:*** A group of participants from University of Cincinnati Science Policy Chapter [wrote about the course in their own words](https://ucsciencepolicy.wordpress.com/2021/02/01/uc-scipol-members-pitch-in-about-what-they-gained-from-the-science-policy-and-advocacy-course-for-stem-scientists/) stating the benefits of the program for their own policy training and careers. Other examples include [Chia Chun Angela Liang](https://www.linkedin.com/in/chia-chun-angela-liang/), an earlier program graduate who founded the [UCI SPAN group](https://www.linkedin.com/in/uci-span-a5a0a921a/) as a result of her growing interest in the field due to this program, and later became a program co-coordinator in 2023.

## Discussion and Future Directions

While limited in timeframe and participation, this certificate program model is unique and has proven successful and could be further expanded to other universities across the country and internationally, in order to support trainees who may have less opportunities. Increasingly, there is also a growing interest from more experienced professionals to continue honing their training, and similar programs could also potentially be developed for other career stages beyond early career.

> ***Reducing Barriers*:** We wanted to ensure that anyone interested in science policy from around the world could enroll in the program. We instituted an application process after the first cohort in order to limit to folks who were really interested in policy. We also focused to an extent on students with fewer opportunities. We addressed barriers to access for science policy and advocacy opportunities by taking into account location (including US states that traditionally had fewer such training opportunities) and not placing any restrictions including for trainees outside the US. In addition to eliminating citizenship requirements - including for international trainees who often have fewer such opportunities to take advantage of US opportunities - we also included undergraduate students and postdoctoral researchers in the program. While several programs exist that cater to graduate students, other populations prior to or post graduate school are often excluded from these opportunities.
>
> ***Format and Cost*:** Provided that the program was offered online, it helped break the challenges of attending in-person networking events and removed geographical barriers for trainees to participate. In addition, the program was offered free of charge, which may have helped remove barriers for trainees from different socio-economic backgrounds. The only technological requirement for the program was to have a computer with stable internet connection for watching publicly available online videos and attending flipped classroom format activities during program sessions. Given pandemic constraints, we were lenient when it came to providing submission assignment extensions beyond the deadline for those who needed or requested it, and on a case-by-case basis allowed participants who could not finish the program the first time due to being sick with COVID-19 or other reasons to return and graduate from the program in subsequent years.
>
> ***Workforce Development:*** The ability to break barriers and silos for groups of trainees that don’t often have opportunities was an important part of the program. Without voices that represent the wide-ranging needs of stakeholders invested in societal change, the lack of participation in the science policy workforce constrains the resulting policymaking process to benefit a narrower subset of society and only allows a small segment of the population to contribute. Evidence-based science policy is thereby critical for maintaining the competitiveness of the research enterprise in the US, and the science policy workforce needs to reflect the people it serves. As policy change impacts every individual in our society, it is imperative to ground the policymaking process in diverse and inclusive viewpoints and backgrounds. The science policy workforce - which we define as professionals who transitioned into a science policy career from a variety of backgrounds and career stages - is not exempt from this principle, when it comes to researchers interested in STEM fields applying their scientific knowledge and backgrounds to the policymaking process.
>
> ***Policy Skills Gained:*** Overall, for all three cohorts, program participants learned fundamental skills in policymaking through a variety of different types of learning. In addition, many of the program participants went on to pursue a full-time career in science policy through a number of opportunities or transition points (e.g. policy fellowships). The program also provided networking opportunities for participants, which is essential for learning about the field and building avenues of professional development. The program had great success in all three years, and several program participants provided testimonials about how it helped their careers. We hope this program will start to shift the paradigm that anyone can play a role in policy, that barriers should be eliminated and that we will see such similar opportunities in the future.

## Program Resources

The organizers created a [YouTube channel](https://www.youtube.com/channel/UC38Ptj2Fj0diLsG2Y4M513Q) dedicated to this program, where all the module videos from all three years are open source for anyone to access and learn from (**Supplementary Figure 2**, **Table 3**). We believe that this mechanism will help trainees who do not have time or are otherwise unable to attend this highly engaging program but can watch videos and learn about the policy world in a more passive manner. Importantly, this channel will also serve as a resource for those who attended the course and would like to go back and refresh their policy training by watching online videos. Another use for these YouTube videos is to help other institutions adapt the program content to their liking while building similar programming catering to their PhD students and postdocs.

As a result of the program, Adriana Bankston and Harinder Singh who originally spearheaded the program, later became [ARIS fellows in the 2022-2023 cohort to develop the science policy workforce](https://researchinsociety.org/project/developing-next-gen-workforce-science-policy-bridge-science-society/) based on these teachings. To facilitate broader exposure of this program and recapitulation at other institutions, they developed a toolkit for universities to create similar training by which to teach policy skills which might also fill the gap in training for PhD students and postdocs applying for science policy fellowships with limited or no exposure or experience (15).

The toolkit is meant to be utilized by professionals involved in career and professional development, such as university administrators and teaching faculty. Thus far, toolkit elements include an [InterSECT Job Sim on the science policy course](https://intersectjobsims.com/library/science-policy-course/) and a [Padlet toolkit](https://padlet.com/scipolucirvine/tool-kit-for-engaging-stem-scientists-in-science-policy-89uq2s3bsp5lsx31) based on the curriculum from this program, with video module links posted in the Padlet boxes. Access to the Padlet link is free and can be customized for any program, and those desiring to design a similar program can copy and paste the video modules in the module sequence of their choice. These tools, along with the [NPA slide presentation](https://www.youtube.com/playlist?list=PLxh_oLbA4CUGWn6Xf6wWH9StmBUTQ81pY) for the program will help university training professionals to create a general syllabus for a full virtual course similarly to this one, or develop their own variations of the program to suit their needs and desired outcomes through the ability to pick and choose modules and design different versions of the program.

## Community Building

The program was offered in a very minimalistic budget for the organizers because we did not invite policy trainers who required high level reimbursements or compensations for their time. Instead, we utilized the network of policy professions from the organizations participating in the program to recruit speakers and reviewers. The alumni and guests in the program were happy to offer their time and excited to share their insights from the field with the program participants, stating multiple times how happy they were that such a program was created.

To ensure the sustainability of the program, we used this program to also build a “train-the-trainer” model, where student volunteers were able to participate in organizational aspects of the program as co-coordinators, and we brought them back the following year to serve as speakers and judges for the elevator pitch competition. We believe this is a helpful practice for continuing to build the field by developing future leaders who can utilize gained skill sets for future cohorts.

In addition, program alumni from the first cohort wanted to give back by serving as speakers, panelists, and volunteering for networking sessions, and were invited back in subsequent years. This model has been very successful and is ensuring the low budget sustenance of the policy program. In the future, we plan to engage more program alumni at multiple levels within the design of the program, in order to continue developing the next generation of policy professionals and foster their continued engagement in the program and with the field of policy and advocacy.

Another example of how trainees were exposed to the field early on was through the opportunity to listen to and interview professionals for the GPS-STEM Radio - Science Policy Podcast, facilitating networking and integration into the field in a low key manner that was mutually beneficial. These opportunities were also fostered through a closed program [LinkedIn group](https://www.linkedin.com/groups/13880099/) and private Slack channel for each class year, in order to build a community of policy and advocacy enthusiasts and provide an opportunity for program participants, alumni, speakers and organizers to connect to each other. In the future, we plan to develop a more extensive alumni programming using prior speakers and participants who would like to mentor participants in subsequent program years.

## Conclusion

We hope this program and our findings shed some light onto what is possible to achieve in policy training through a strong dedication of program directors, coordinators, speakers, reviewers and program participants, as well as a strong collaborations between universities and other stakeholders who were instrumental in designing and sustaining this program for three years, and who are heavily invested in building future leaders to drive policymaking at all levels both in the US and internationally.

## Acknowledgements

We would like to thank program coordinators (Ria Desphande, Klebea Sohn, Melania Abrahamian, Pawan Upadhyay and Chia-Ching Liang), founding partners (Melissa Varga, Steve Allison) and the numerous speakers, reviewers, and trainers for participating in the program for their contributions. We also thank our program partners and supporters for their help and support with financial contributions and program dissemination to recruit top applicants to the program and ensure their success. For questions about the program or if you’d like to get involved, please contact the corresponding author Adriana Bankston at.

## Supplementary Figures

**Supplementary Figure 1.**
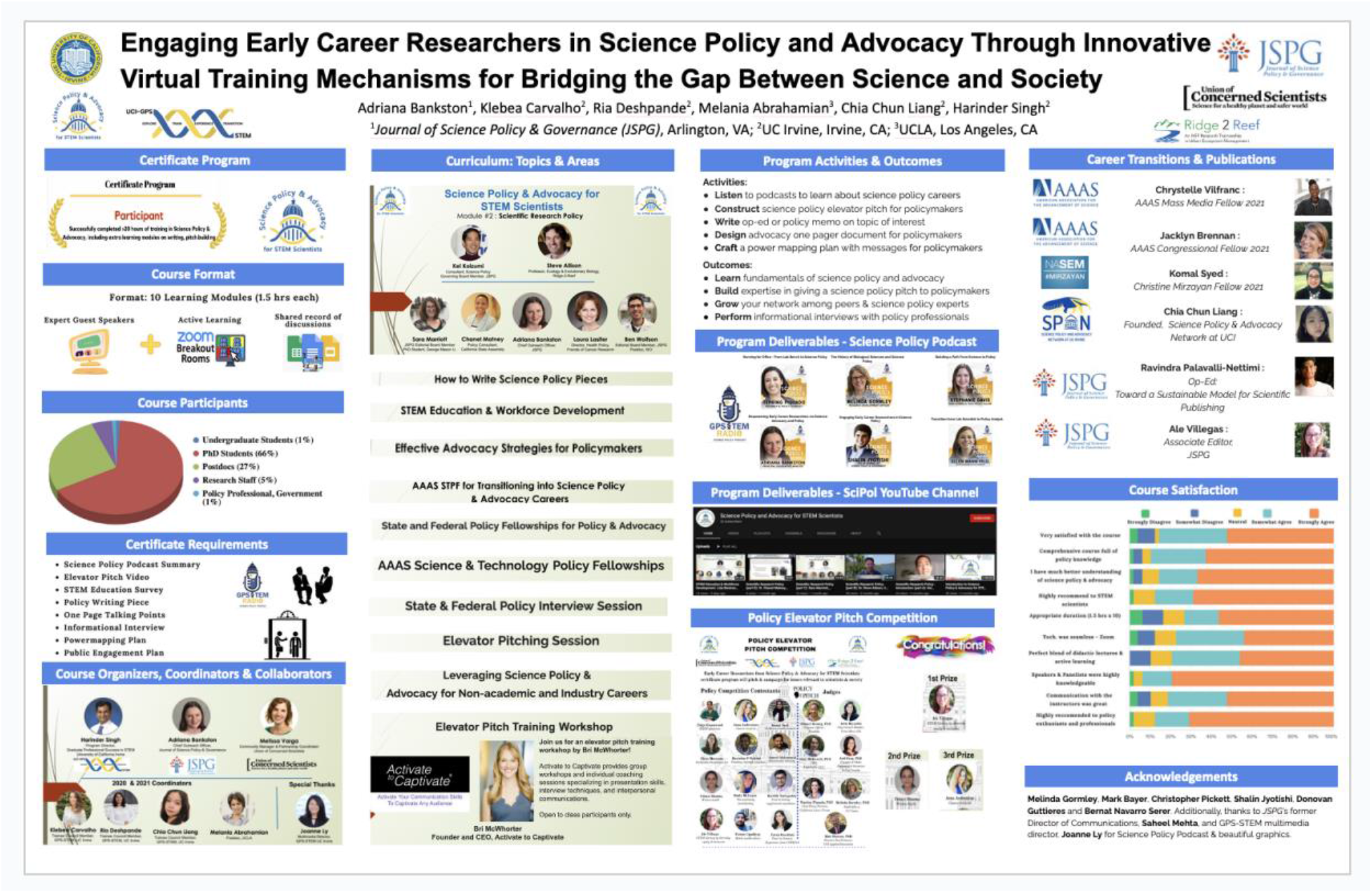
National Postdoctoral Association Poster on Certificate Program. This poster was presented at the National Postdoctoral Association annual meeting in 2022 prior to the third cohort. <u>Poster slides Poster video</u>

**Supplementary Figure 2.**
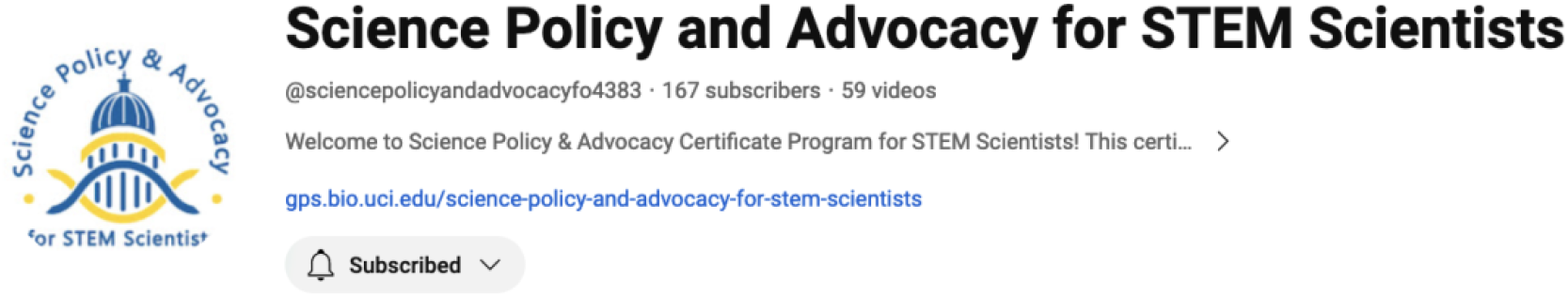
YouTube Videos from the Three Program Cohorts. YouTube videos from the 2020 program are linked in Table 3 with <u>YouTube videos from</u> <u>2021 program</u> and <u>YouTube videos from 2023 program</u> in this YouTube channel.

## Supplementary Tables

**Supplementary Table 1.** Number of Program Participants. Registrants are trainees who expressed interest in and signed up to enroll in the course in all 3 years, out of which a select number were chosen from the application process in 2021 and 2023. *Application process instituted for the 2021 program and also used for 2023.

| Year | Registrants | Offered | Graduated |
| --- | --- | --- | --- |
| 2020 | 135 | 135 | 52 |
| *2021 | 158 | 100 | 48 |
| *2023 | 210 | 120 | 70 |

**Supplementary Table 2.** Career Stage for Program Participants. These data constitute the number of trainees who enrolled in the program initially from PhD students, post-PhD, and pre-PhD.

| Year | PhD Students | Post-PhD (Postdoc, Research Staff, Project Scientist, Specialist) | Pre-PhD (Undergraduate, Masters) |
| --- | --- | --- | --- |
| 2020 | 129 | 34 | 3 |
| 2021 | 134 | 19 | 5 |
| 2023 | 94 | 57 | 39 |

**Supplementary Table 3.** Location for Program Participants. These data constitute the number of trainees located in the US or abroad while enrolled in the program, based on their affiliation at the time of enrollment.

| Year | Based in the US | Non-US (Located abroad) |
| --- | --- | --- |
| 2020 | 121 | 14 |
| 2021 | 134 | 24 |
| 2023 | 114 | 96 |

